# Research productivity and collaboration of the NIH-funded HIV Vaccine Trials Network: a bibliometric analysis

**DOI:** 10.1101/2020.06.05.136846

**Authors:** Jonathan Nye, M.Patricia D’Souza, Dale Hu, Dolan Ghosh

**Affiliations:** Office of the Director, NIAID, NIH, Bethesda, Maryland, USA; Division of AIDS, NIAID, NIH, Bethesda, Maryland, USA

**Keywords:** Bibliometrics, Outcomes, HIV vaccine, HIV

## Abstract

**Objectives:** To assess the scientific productivity and impact of the HIV Vaccine Trials Network (HVTN) over the last two decades and to examine how research collaboration has evolved over this time in the HIV vaccine field.

**Design:** This section does not apply since this is a bibliometric study.

**Methods:** A systematic bibliometric analysis was conducted to identify all HIV vaccine and HVTN associated publications from 1999-2019. All publications were sourced from the NLM Pubmed database and funding information was obtained from the SPIRES and iSearch databases. Both the InCites and iCite databases were utilized for impact metrics. Finally, HVTN clinical trials were obtained from clinicaltrials.gov. Multiple field normalized citation metrics, such as the relative citation ratio (RCR) and number of publications in the top 1% and 10% within their respective field, were used to gauge scientific impact of publications. Network analyses were used to examine collaboration among the most prolific researchers in the HIV vaccine research.

**Setting:** This section does not apply since this is a bibliometric study.

**Participants:** This section does not apply since this is a bibliometric study.

**Intervention:** This section does not apply since this is a bibliometric study.

**Results:** 79 clinical trials were funded by the HVTN from 1999 to 2019. These were carried out via a network of trial sites in 23 countries and 94 cities around the world. In total, 465 publications (89.5% original research articles, 7.3% reviews, and 3.2% other) acknowledged funding from the HVTN. Impact analyses using multiple field normalized metrics revealed that HVTN publications are highly cited with a mean RCR of 1.8. 10,481 HIV vaccine related publications were used to analyze collaboration in this field. Compared to the field as a whole, publications attributed to the HVTN had significantly more authors per publication (p-value < 0.001) and our network analysis found that HVTN-associated authors also had a higher degree (p-value < 0.01).

**Conclusions:** Bibliometric analysis of the last two decades of HIV vaccine research by the HVTN revealed that in addition to conducting a large number of clinical trials worldwide, the network produced high impact publications and was associated with increased collaboration among researchers.

**ARTICLE SUMMARY:** *Strengths and limitations of this study:*

- To the best of our knowledge, this is the only study that has provided a systematic bibliometric analysis of the HVTN since its inception.
- Studies like this can illustrate overall outcomes of large clinical network programs.
- Advanced field normalized metrics were used to provide the most accurate measures of productivity, impact, and collaboration.
- Identification of HVTN publications using funding acknowledgements can lead to an underestimate of the number research articles since not all research articles include grant funding information.
- Variations in a single author’s name across publications can make name disambiguation difficult when performing network analyses.

## INTRODUCTION

Human immunodeficiency virus (HIV) remains a public health concern with an estimated global prevalence of 36.9 million HIV-infected persons worldwide and 1.8 million new infections per year ^1^. Remarkable progress in treating HIV/AIDS has been made after almost four decades of active research since the first cases of AIDS were reported. Prevention and treatment have dramatically improved as a result of increased testing and treatment with anti-retroviral therapy (ART) ^2^. However, there are still no licensed vaccines to prevent HIV infection, even though a vaccine will likely be essential to achieve a long-lasting end to the global pandemic ^3^.

Several HIV vaccine efficacy trials were conducted between 2004 and 2009. One of these trials, known as RV144, resulted in the first vaccine regimen to exhibit a protective effect, suggesting that an effective vaccine might be achievable ^4^. Since then, researchers around the world have worked to build on these findings in hopes of developing a more effective and durable immune response capable of preventing HIV infection. The National Institutes of Health (NIH) funds the large majority of research on HIV/AIDS vaccines in the world. Indeed in 2018, 85% of funding for HIV vaccine research worldwide was contributed by only two major funders, the NIH and the Bill and Melinda Gates Foundation ^5^. Within the NIH, one institute in particular, the National Institute of Allergy and Infectious Diseases (NIAID), through its Division of AIDS (DAIDS) has led the effort to develop a safe and effective vaccine and has supported a robust body of HIV vaccine-related research from preclinical and translational research to clinical trials. In addition, it has established and supported several large networks dedicated to conducting HIV/AIDS clinical trials both within the United States and globally.

Since it was established in 1999, the NIAID supported HIV Vaccine Trials Network has conducted the majority of clinical trials of preventive HIV vaccines worldwide ^6^. The HVTN is comprised of an international group of scientists, educators, and community members whose mission is to support the development of a safe and effective vaccine for prevention of HIV infections. It conducts all phases of clinical trials, from testing safety and immunogenicity of vaccine candidates to evaluating vaccine efficacy. It is made up of three parts: the Laboratory Center, the Statistical and Data Management Center, and the Leadership and Operations Center ^7^. All three of these work closely with the clinical research and trial sites. As the federal government funder of non-governmental networks like the HVTN, DAIDS plays a major collaborative role as not only the funder but in scientific and protocol development, trial and safety monitoring, laboratory and other support, in addition to serving as the regulatory sponsor. The HVTN’s trial sites are located at research institutions around the world while the vaccine products come from various developers, both for profit and academic investigators. This structure allows it to streamline HIV vaccine testing and to reach populations severely impacted by the HIV/AIDS epidemic in both the U.S. and abroad ^8^.

Although the HVTN is one of the largest and longest lasting HIV research programs, its productivity and impact has not been well-documented in the literature. While previous studies have examined research outputs ^9^, expansion of subject areas ^10^, collaborations ^11^, and the geographic distribution of HIV research ^12^, they have been relatively limited in scope in terms of geographic region or time ^13–16^. Moreover, despite the growing importance of scientific collaborations ^17^, studies examining collaborations within HIV clinical trials networks have been limited to only a few years ^11^. Previous work has outlined the scientific achievements over the first decade of the HVTN however a bibliometric analysis of the program has yet to be done ^7^.

Our study seeks to build on previous work by providing a comprehensive bibliometric analysis of the HVTN from 1999-2019, including an overview of the international network of clinical trial sites utilized by the program and an in-depth examination research outputs such as clinical trials and number of publications in combination with advanced field normalized metrics to assess the impact of this work. We also show how collaboration has evolved in the HIV vaccine field as a whole as well as among HVTN investigators. Together, this work provides an overview of the productivity and impact of the HVTN since it was first established 20 years ago.

## METHODS

Both publicly available and internal NIH databases were used to gather data for the study. All analyses and visualization were carried out using the R programming language.

### HVTN clinical trials

A comprehensive list of clinical trials that were attributed to the HVTN was obtained from the ClinicalTrials.gov database through 2019 by searching for trials with keyword “HVTN”. From this list we kept only trials with an HVTN identifier listed in the Acronym or Other Study ID. Finally, this list of trials was manually curated by program staff. This resulted in a final list of 79 clinical trials.

### Geographic distribution

Data on HIV prevalence among people ages 15-49 in 2017 was obtained from the World Health Organization. HVTN clinical trial sites were retrieved from ClinicalTrials.gov. Out of the 79 total clinical trials 77 had location information associated with them. Counties and states comprising >50% of new HIV infections were obtained from the Ending the HIV Epidemic initiative. Mapping of global clinical trial sites and HIV prevalence was done using the ggplot2 package.

### HVTN publications

The iSearch platform is a suite of tools available to NIH staff that provides access to a comprehensive, curated, extensively linked data set of global grants, patents, publications, clinical trials, and FDA-approved drugs. The iSearch Publications tool utilizes the NLM PubMed and SPIRES databases. The SPIRES database contains positively, verifiable mappings between scientific publications and NIH grant numbers and is available to NIH staff. Using the iSearch Publications tool, we searched for all publications that acknowledged HVTN grant funding using grant numbers. NIH’s publicly available iCite tool ^18^ was used to distinguish research articles from derivative or non-research articles. The iCite article type classification is based on PubMed “Publication Type” tags. Of the 465 publications citing HVTN support, 416 research articles were retained for analysis. The other 49 publications were review articles, commentary, or other non-research articles. The HVTN publications found in the SPIRES system include only those publications that cite support from NIAID funding. Not all publications contain such citations.

### HIV vaccine publications and coauthorship network analysis

iSearch was also used to identify a larger set of publications encompassing the HIV vaccine field, using the following search terms applied to publication titles and abstracts: (HIV* AND VACCIN*), (AIDS AND VACCIN*), (Antibodies AND Neutralizing AND (HIV* OR AIDS)). This resulted in 16,643 publications published between 2000 and 2019. From this dataset, 12,426 were identified as research articles using the iCite article type classification described above. In addition, to enrich for articles that were specific to the HIV vaccine field, we excluded articles where HIV/AIDS was not the primary focus of the article. Therefore, we removed publications that contained the following keywords or parts of keywords in the title: tuberculosis, hepatitis, influenza, papilloma, pneumococc, meningococc, herpes, streptococc, HBV, HCV, or yellow fever. This left us with a final list of 10,481 HIV vaccine research articles including 281 acknowledging HVTN funding.

From the HIV vaccine publication dataset, we created undirected and unweighted coauthorship networks using the igraph and ggplot2 packages in R. We built two coauthor networks, one spanning the years 2000-2009 and another from 2010-2019 to capture how collaboration in the field has changed over time. The network layouts were generated using the Kamada-Kawai force-directed algorithm ^19^. Authors publishing under a number of different name variations is a challenge in creating coauthor networks, so to ensure the quality of our results we used a custom script for author disambiguation. Briefly, it calculates the number of name variations among authors with the same last name and first initial combination to determine how many possible variants there are for each combination. Authors were separated into categories of high or low confidence based on this value. Low confidence combinations with many different names were manually corrected and high confidence combinations having little or no variation in naming were automatically corrected. For the sake of simplicity and clarity, only the top 150 most prolific authors in each time period were used in our networks. This allowed us to see how collaboration among the most prolific investigators in the HIV vaccine field evolved while avoiding overly dense and crowded networks. Furthermore, we identified all of the investigators on publications acknowledging HVTN grant funding and highlighted nodes representing these HVTN associated investigators and the edges connecting them.

## RESULTS

### HVTN clinical trials

Of the 79 trials funded by the HVTN through 2019, 61 were Phase I, 6 were Phase I/Phase II, 10 were Phase II, 1 was Phase II/Phase III, and 1 was a Phase III trial (Fig 1). In total, over 26,000 participants were enrolled over this time period. The largest portion coming from the large proof-of-concept and efficacy trials which enrolled 18,658 participants. The smaller Phase I and II trials testing safety and immunogenicity enrolled 7,978 participants.

**Fig 1.**
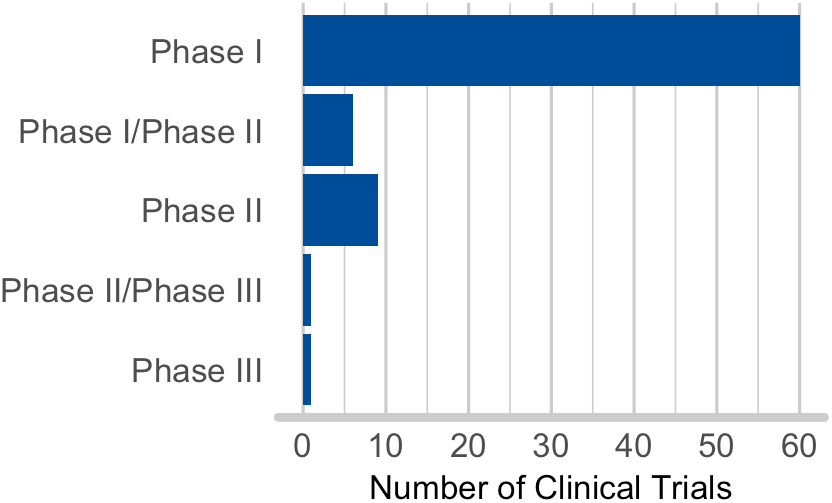
HVTN clinical trials. Number and phases of clinical trials conducted by the HVTN up until 2019.

### Geographic distribution

The global network of all HVTN clinical trial sites up until 2019 and the prevalence of HIV among people ages 15-49 in 2017 is shown in Figure 2. The HVTN had clinical trial sites in 23 countries and 94 cities worldwide. Of the 79 trials, 65 had a US component. These were carried out in 20 different states and 31 cities around the country. Many of these clinical trials took place in communities that have been most affected by the HIV epidemic. Recently, 48 counties and 7 states have been identified that account for >50% of all new HIV diagnoses in the United States between 2016 and 2017. State and county level data indicate that, the HVTN has had trials in 21 of these counties and 2 states.

**Fig 2.**
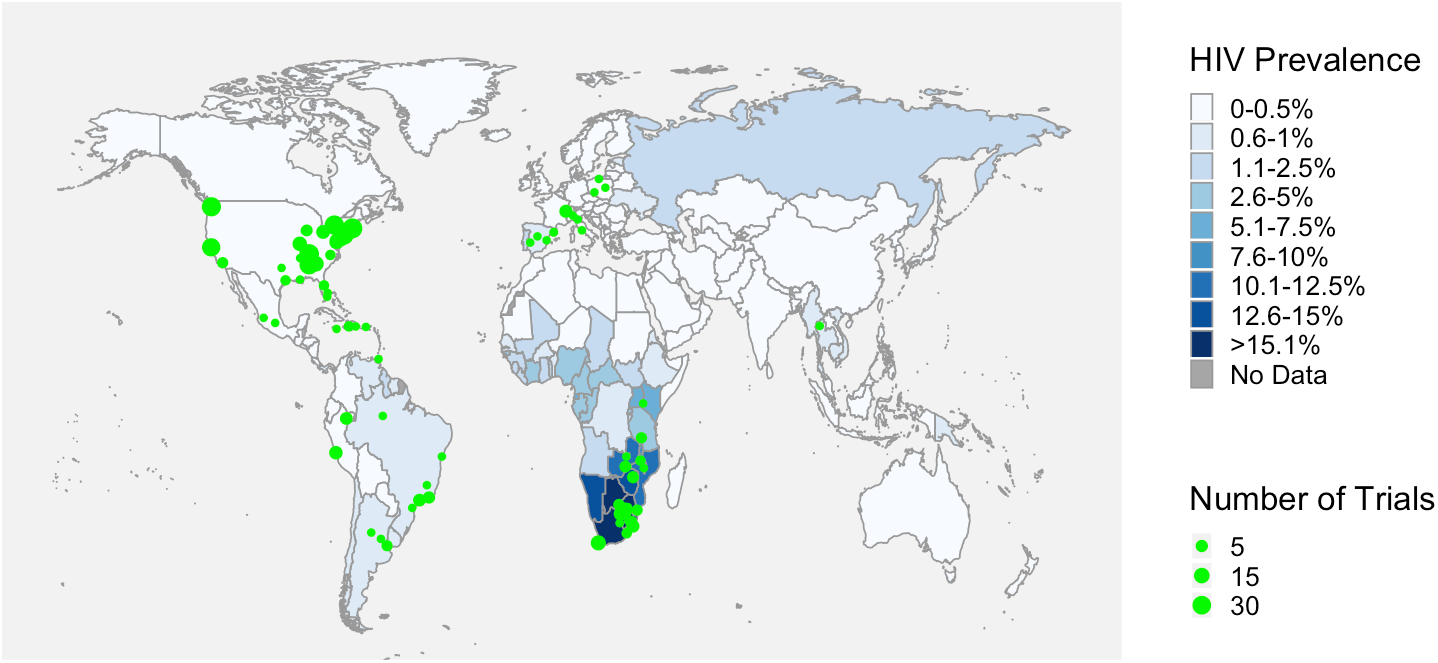
HVTN clinical trial sites. HIV prevalence among people ages 15-49 in 2017 and HVTN clinical trial sites around the world.

### HVTN productivity and impact

To get a better understanding of the impact and productivity of the HVTN we identified all publications that acknowledged grant support from 1999-2019 (Fig 3). In total there were 465 publications (89.5% original research articles, 7.3% reviews, and 3.2% other). Out of the entire set of publications we decided to focus specifically on the 416 original research articles when assessing the performance and impact of the HVTN since these are the best indicators of the network’s scientific contributions. Analysis of the number of research articles per year revealed a dramatic increase in publications per year beginning in 2011. After reaching a peak of 50 publications in 2014 and again in 2016 the number of publications declined to 32 research articles in 2018. However, this trend was reversed in 2019 when 42 articles were produced by the network.

**Fig 3.**
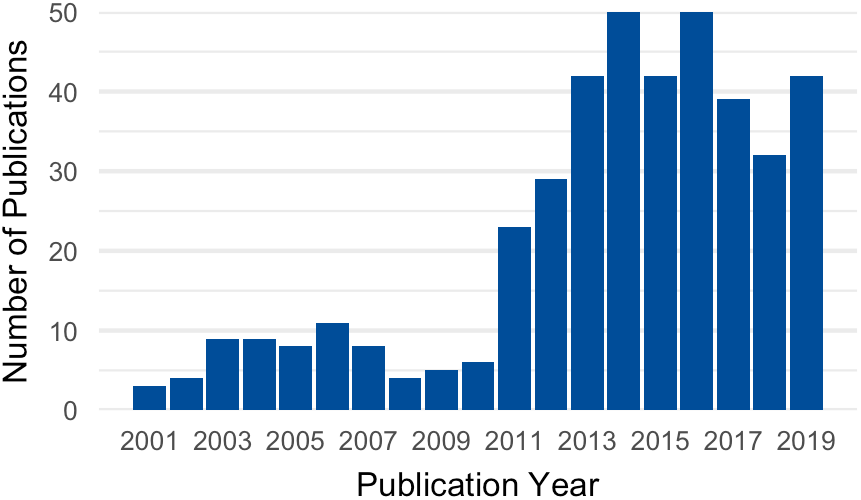
HVTN productivity and impact. Number of research articles per year acknowledging support from the HVTN from 1999-2019.

In addition to the productivity, we wanted to gauge the performance and impact of these publications using citation-based metrics including multiple field normalized indicators (Table 1). We found that HVTN research articles had been cited a total of 12,521 times with a median number of citations per paper of 10 and a mean of 30.1 (range 0-1088). In addition, 99.1% were cited at least once after 5 years compared to 88% of all articles in iSearch database. In order to determine how these publications performed relative to their field of study we used two separate metrics. The first is the relative citation ratio (RCR) which uses a novel method based on the paper’s co-citation network to provide a field normalized metric ^18^. We found that HVTN publications had a mean RCR of 1.8 (range 0-47.4) meaning that on average they were cited 1.8 times more than expected. In addition, we used the InCites database which organizes research articles by publication year and subject area based on journal category to analyze the percentile rank of 399 HVTN publications found in their collection ^20^. We found that after normalizing for time and subject area 22.1% were in the top 10% most cited papers indicating that these publications were represented more than twice as much as expected in this highly cited category within their respective field. Moreover, we found that 5% of these publications were in the top 1% most cited papers, meaning that HVTN supported papers were represented 5 times as much compared to other papers in their field and is better than all NIH supported papers that made the top 1% cited tier ^21^.

**Table 1.**
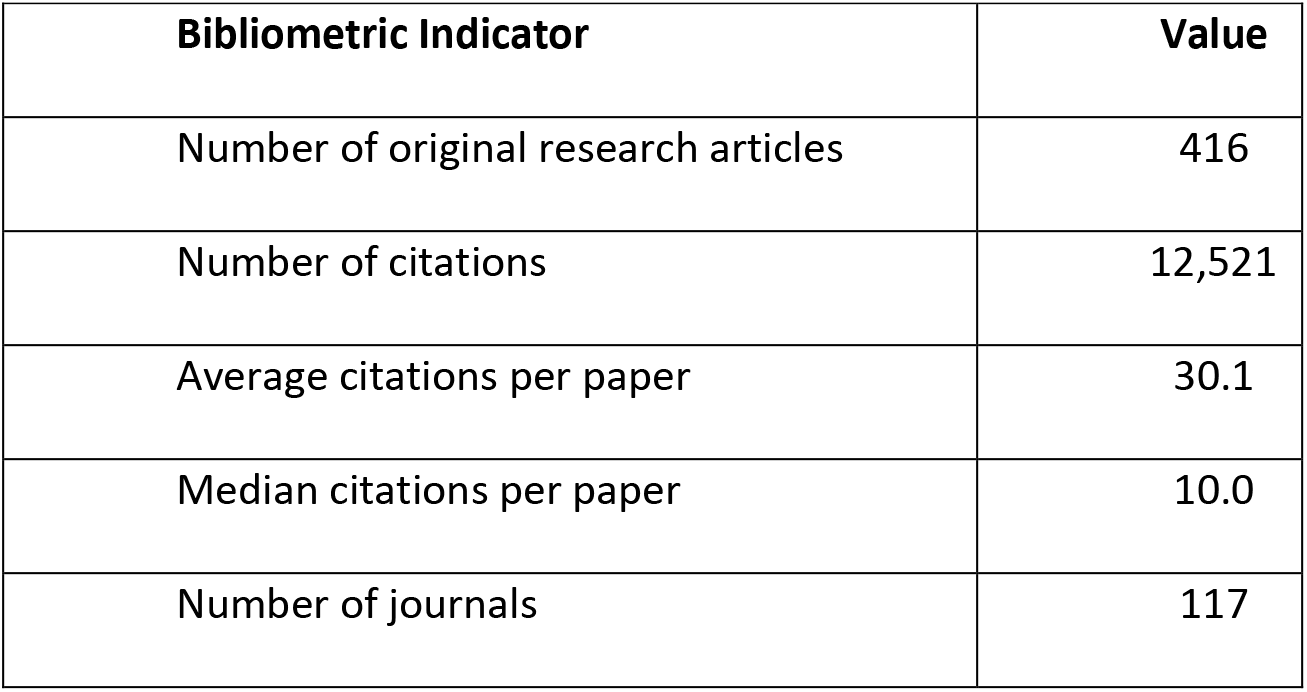
Summary of bibliometric indicators.

| **Bibliometric Indicator** | **Value** |
| --- | --- |
| Number of original research articles | 416 |
| Number of citations | 12,521 |
| Average citations per paper | 30.1 |
| Median citations per paper | 10.0 |
| Number of journals | 117 |

To compare the HVTN’s productivity with the HIV vaccine field we utilized a keyword search to identify 10,481 unique publications that represent the HIV vaccine field from 2000-2019. To analyze how the field changed over time we looked at two different time periods, one spanning the years 2000-2009 and another from 2010-2019. Our analysis revealed that productivity of the HVTN increased from 44 publications in time period 1 to 237 publications in time period 2. During this timeframe the HIV vaccine field grew with the total number of publications increasing from 4,471 in the earlier time period to 6,010. HVTN publications had a 5.4-fold increase from the first time period to the next while the HIV vaccine field grew by 1.3-fold and the iSearch index grew by 1.7-fold during the same time periods. In comparison, HVTN associated authors had a mean number of publications of 28.9 from 2000-2009 and 60.4 from 2010-2019. On average, the authors on papers acknowledging HVTN funding published slightly more often and showed a greater increase in productivity between the two time periods compared to the top 150 most prolific authors in the field.

### Collaboration in the HVTN

As a measure of collaboration, we looked at the number of authors per publication between these two time periods. We found that the mean number of authors on HIV vaccine publications increased significantly (Welch’s two sample t test, df = 10,421, p-value < 0.001) from 7.5 (range 1-52) in the first time period to 9.4 (range 1-61) in the second time period. Similarly, HVTN publications in this dataset also tended to have significantly (Welch’s two sample t test, df = 127.6, p-value < 0.001) more coauthors with the mean number of authors rising from 8.8 (range 1-21) to 14.4 (range 1-56) respectively. When we compared number of authors on HVTN publications to non-HVTN publications we found that HVTN publications had a significantly higher (Welch’s two sample t test, df = 244.1, p-value < 0.001) number of authors per paper during the second time period suggesting that HVTN authors collaborated more compared to authors in the HIV vaccine field. As an additional measure of collaboration, we created two coauthor networks one spanning each of the two different time periods (Fig 4). These networks have been shown to be a very useful tool for the analysis of collaboration within a field ^22 23^. Each node in the network represents an author while a connection (edge) between these nodes indicates coauthorship. We restricted the networks to the top 150 most prolific authors in each time period. These authors were responsible for greater than a third of the publications and were the most connected. Author names were disambiguated using a script described above and further checked manually to ensure accuracy. Edges connecting authors on HVTN publications have been highlighted. Next, we calculated the degree of each investigator which corresponds to the number of authors that individual has published with and is equal to the number of edges in the network for that person. The network analysis revealed that collaboration increased over time in the HIV vaccine field with the average degree rising from 19.4 between 2000 and 2009 to 63 in the following years. In addition, we examined HVTN associated authors and found a similar trend in which the mean degree increased from 27.6 and 66. This was significantly higher than for non-HVTN authors (Welch’s two sample t test, p-value < 0.01), mean of 17.5 from 2000-2009 and mean of 52.1 from 2010-2019. Finally, our analysis revealed that the number of the HVTN associated investigators represented in the networks more than tripled over these two time periods, increasing from 28 to 116 individuals.

**Fig 4.**
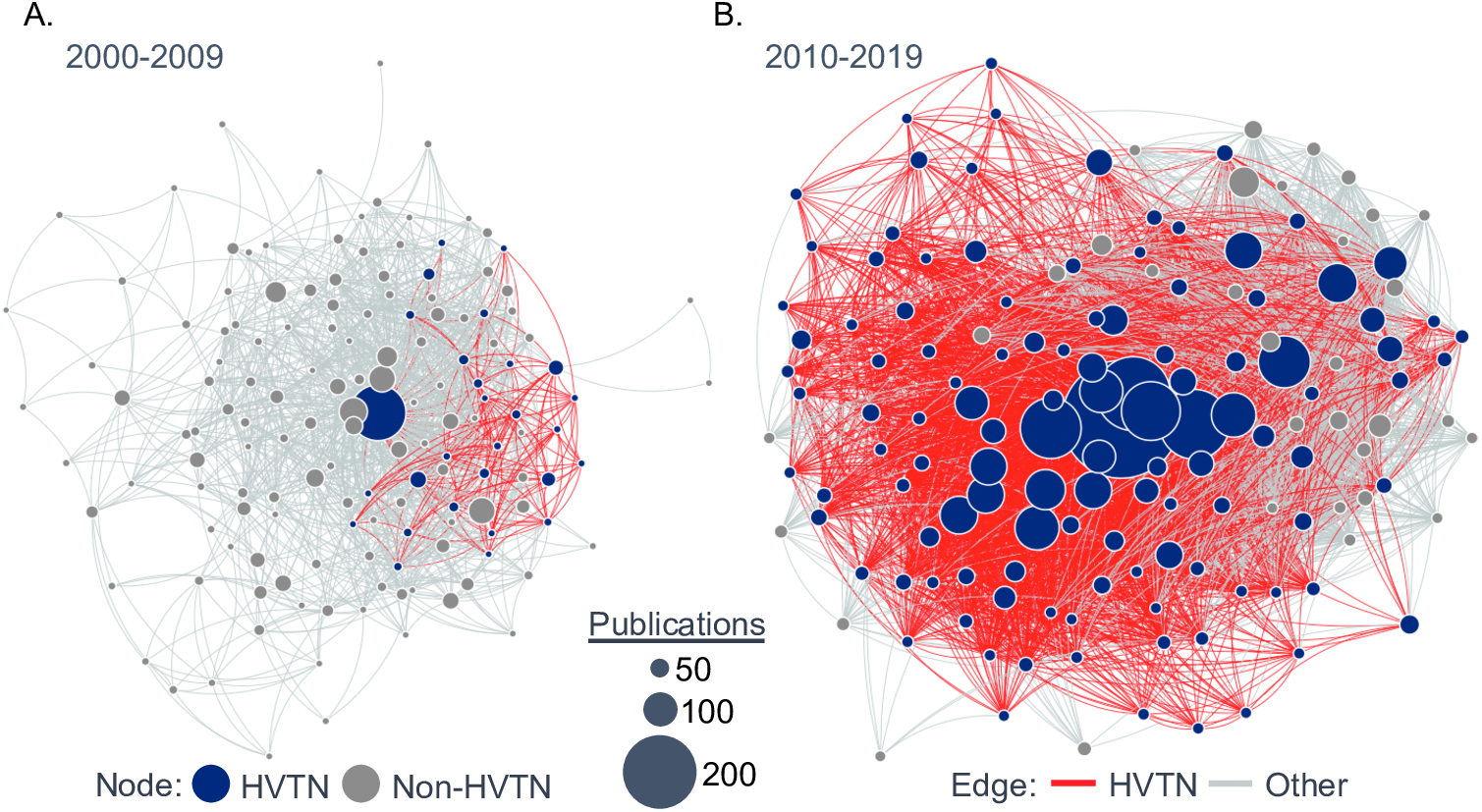
HIV vaccine coauthor networks. Each node represents one of the top 150 most prolific authors in the HIV vaccine field from either (A) 2000-2009 or (B) 2010-2019. Node size indicates the total number of publications per author. Edges connecting nodes indicate coauthorship. Edges connecting authors on HVTN publications are highlighted.

## DISCUSSION

The development of a safe and effective HIV vaccine is entering a very exciting phase with four efficacy trials underway, more than any other time in the history of HIV vaccine development. These developments represent the culmination of many years of preclinical research and clinical trials, with most of this research funded by NIH making this a perfect time for assessing the HVTN program. Clinical sites are an essential component of the network, therefore, it must support a robust global network capable of handling a large number of clinical trials. Indeed, we find that the HVTN supports clinical sites in 23 countries and 94 cities worldwide. Many of these countries have been the hardest hit by the AIDS epidemic including South Africa, Zimbabwe, and Botswana among others. The global expansion of trial sites has coincided with the shift in HIV vaccine antigen design. Between 1985 and 2005, over 90% of candidate HIV vaccines were clade B based antigens, while in the last 10 years, >80% of the HVTN portfolio involves clade C and mosaic envelope antigens ^24–26^. Our studies indicate that the HVTN is furthering its program goals of reaching populations severely impacted by the HIV/AIDS epidemic in both the U.S. and abroad ^8^.

In addition to carrying out a large number of vaccine clinical trials, the HVTN in general has increased productivity over time, publishing more since 2011. This increase in publications may be due in part to new insights gained from the RV144 trial in which the first partially effective HIV vaccine was tested. Furthermore, we found that HVTN publications were high impact as shown by multiple field normalized citation metrics including RCR and the percentage of publications in the top 1% or 10% in their respective field of study. Many of these articles summarize the major accomplishments throughout the life of the network such as the development and analyses of numerous new vaccine approaches, products, and adjuvants ^26–33^. Additionally, our analyses revealed that HVTN associated PIs more than tripled among the top 150 most prolific PIs in the field from the first time period to the next.

Our analysis of research articles in the HIV vaccine field revealed that collaboration increased significantly during the assessed time period as indicated by an increase in the mean number of authors per publication. Moreover, this increase was even higher for HVTN associated investigators compared to the field. Our coauthor network analysis of the top 150 most prolific authors showed that collaboration among them also increased substantially from 2000 to 2019 as indicated by the tripling of the average degree. In addition, we found that HVTN associated investigators had a significantly higher degree compared to non-HVTN investigators. This difference was likely driven in part by increased publication frequency but also by larger team sizes. Thus, the HVTN’s unique structure may create an environment that fosters collaborations to stimulate interdisciplinary clinical research.

Scientific research collaboration is critically important in a complex and multidisciplinary field such as HIV vaccine development as it allows improved sharing of knowledge and expertise as well as the pooling of resources and data. Increasingly sophisticated technologies and the massive amounts of data that is being generated means that more and more researchers must specialize and focus their resources. In turn, increasing specialization of research scientists means that successful research requires increasingly larger, multidisciplinary collaborations and sharing of knowledge. This trend was documented across many disciplines including science and engineering, but it is certainly true for as specialized a field as HIV vaccine development ^17^. Therefore, HVTN’s focus on data sharing and collaboration may help researchers to capitalize on the knowledge gained from its different teams to carry out multidimensional analyses.

Beyond the productivity, influence, and impact measured in this study, the NIH values work that culminates in advances to human health, a process that historically takes decades. Insights into how to accelerate this process may come from quantitative analysis. Metrics have facilitated quantitation of the diffusion of knowledge from basic research toward human health studies, by examining the type rather than the count of citing articles. Insights into how to accelerate this process will probably come from quantitative analysis ^34^. Comprehensive evaluation programs will need to incorporate these additional metrics that can capture other types of outcomes such as the value of innovation, clinical outcomes, novel vaccine platforms, research enabling vulnerable populations, global collaborations, and training the next generation of scientists

## Acknowledgements

We would like to thank Mary Marovich, Ariel Zane, Marie Parker and Jim Onken for their helpful guidance and feedback.

## Author contributions

PD, DH, and DG devised the original idea for the article. JN was responsible for data collection and analysis. DG and JN participated in the development of methodology. All authors contributed to drafting and revising the manuscript.

## Data sharing statement

No additional data available.

